# DNA translocase repositions a nucleosome by the lane-switch mechanism

**DOI:** 10.1101/2021.02.15.431322

**Authors:** Fritz Nagae, Giovanni B. Brandani, Shoji Takada, Tsuyoshi Terakawa

## Abstract

Translocases such as DNA/RNA polymerases, replicative helicases, and exonucleases are involved in eukaryotic DNA transcription, replication, and repair. Since eukaryotic genomic DNA wraps around histone core complexes and forms nucleosomes, translocases inevitably encounter nucleosomes. Previous studies have shown that a histone core complex repositions upstream (downstream) when SP6RNA or T7 RNA polymerase (bacterial exonuclease, RecBCD) partially unwraps nucleosomal DNA. However, the molecular mechanism of the downstream repositioning remains unclear. In this study, we identify the lane-shift mechanism for downstream nucleosome repositioning via coarse-grained molecular dynamics simulations, which we validated by restriction enzyme digestion assays and deep sequencing assays. In this mechanism, after a translocase unwraps nucleosomal DNA up to the site proximal to the dyad, the remaining wrapped DNA switches its binding region (lane) to that vacated by the unwrapping, and the downstream DNA rewraps, completing downstream repositioning. This mechanism may have crucial implications for transcription through nucleosomes, histone recycling, and nucleosome remodeling.

**SIGNIFICANCE:** Eukaryotic chromosomes are composed of repeating subunits termed nucleosomes. Thus, proteins that translocate along the chromosome, DNA translocases, inevitably collide with nucleosomes. Previous studies revealed that a translocase repositions a nucleosome upstream or downstream upon their collision. Though the molecular mechanisms of the upstream repositioning have been extensively studied, that of downstream repositioning remains elusive. In this study, we performed coarse-grained molecular dynamics simulations, proposed the lane-shift mechanism for downstream repositioning, and validated this mechanism by restriction enzyme digestion assays and deep sequencing assays. This mechanism has broad implications for how translocases deal with nucleosomes for their functions.

## INTRODUCTION

Nucleosomes are the building blocks of eukaryotic chromosomes and contribute to packaging of genomic DNA into small nuclei. A nucleosome occludes ∼147 base-pairs (bps) of DNA wrapping around a histone core complex from the proteins involved in DNA transactions (DNA replication, transcription, and repair) (1, 2). DNA translocases (e.g., DNA/RNA polymerases, replicative helicases, and exonucleases) unidirectionally move along genomic DNA using nucleotide tri-phosphate hydrolysis energy (3). Translocases frequently move longer distances than the typical lengths of linker DNA (20 to 50 bps) between adjacent nucleosomes without dissociation (4). For example, human RNA polymerase II transcribes DNA by translocating on the ∼2.5 mega-bps dystrophin gene (5). Therefore, translocases inevitably collide with nucleosomes (2, 6, 7).

In a cellular environment, translocases and histone chaperones cooperatively reposition or dismantle histone core complexes through their specific interaction with the complexes (8). The interaction between a translocase with a histone core complex plays important roles in the repositioning. Previous studies proposed that RNA polymerase II (Pol II) repositions nucleosomes by specifically interacting with the histone core complex (by the so-called Pol II-type mechanism) (9–16). In this mechanism, a translocase passes through a nucleosome without changing the histone core complex position by electrostatically interacting with the complex. However, in order to unravel the roles of the interaction between a translocase with a histone core complex during repositioning, a translocase having no specific interaction with the complex serves as a simple model system. In the previous studies, Studitsky and Felsenfeld et al. performed *in vitro* biochemical assays and showed that a histone core complex repositions upstream when SP6 or T7 RNA polymerase partially unwraps DNA wrapping around the complex (9, 17–20). Since RNA polymerase III (Pol III) shows the same behavior, this mechanism is called the Pol III-type mechanism (21). In these assays, a nucleosome position is localized using the property that the DNA fragment occluded by a histone core complex cannot be digested by restriction enzymes. Notably, in these assays, nucleosomes were reconstituted at the end of DNA, precluding downstream repositioning.

On the other hand, Greene et al. performed single-molecule imaging experiments and showed that a histone core complex repositions downstream when a bacterial exonuclease, RecBCD, collides with a nucleosome (6). In this imaging, nucleosomes were reconstituted at random positions on the λ phage genomic DNA, thus not precluding downstream repositioning. However, sub-kbps spatial resolution of the single-molecule imaging prevented them from proposing molecular mechanisms of the downstream repositioning. While Luger et al. proposed the pushing mechanism in which one base-pair unwrapping is coupled to one base-pair downstream repositioning, the molecular detail of this mechanism have not been explored (22).

In this study, we performed coarse-grained molecular dynamics (MD) simulations, proposed the lane-shift mechanism for downstream repositioning, and validated this mechanism by restriction enzyme digestion assays and MNase-seq assays. In this mechanism, after a translocase unwraps nucleosomal DNA up to the site proximal to a dyad, the remaining wrapped DNA switches its binding region (lane) to that vacated by the unwrapping, and subsequently the downstream DNA rewraps, completing downstream repositioning. This mechanism may significantly improve our understanding of the molecular basis of transcription through nucleosomes, histone recycling, and nucleosome remodeling in a more complex cellular environment.

## RESULT

### Simulations of nucleosomal DNA unwrapping by a translocase

We performed coarse-grained MD simulations to investigate repositioning of a histone core complex when a translocase partially unwraps nucleosomal DNA. The simulation system contains the Widom 601 nucleosome-positioning DNA sequence flanked by random 32- and 96-bps DNA sequences upstream and downstream, respectively, together forming a 275-bps DNA (Fig. 1A, Table S1. The DNA bps are numbered relative to the dyad of the 601 sequence). Initially, the histone core complex is formed on the 601 sequence. A model DNA translocase initially loaded on the upstream DNA proceeds in the downstream direction, colliding with the nucleosome.

**Figure 1:**
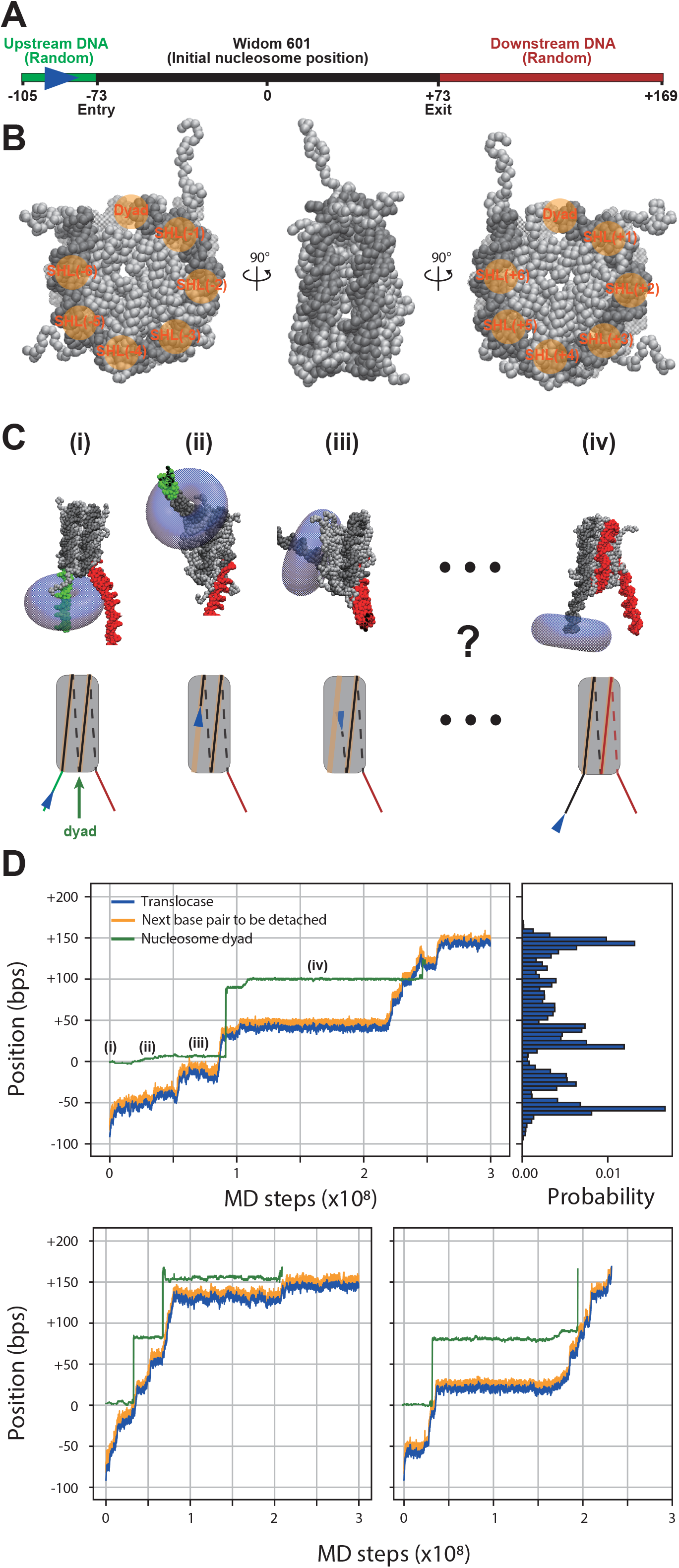
Molecular dynamics simulations of nucleosome unwrapping by a translocase. (A) The map of the DNA sequence used in the simulations. In the initial structure, the Widom 601 sequence DNA wraps around a histone octamer to form a nucleosome. The DNA bps are numbered so that the dyad base-pair position in the initial structure becomes ±0 bps. (B) Coarse-grained model of a nucleosome with the definitions of SHL’s. (C) Snapshots from a simulation and their cartoon representations. The translocase and the histone octamer are colored blue and grey, respectively. The DNA is colored according to the color scheme in (A). (D) Representative simulation trajectories of the positions of the translocase, the base-pair to be detached, and the dyad (top left panel and bottom panels) and a probability distribution of the translocase position (top right panel).

In this simulation, for histones, we used a coarse-grained model in which one particle represents one amino acid (23). A structure-based potential stabilizes the native structure of the histone core complex (24). The interactions between the histones were adjusted so that the complex disassembles in the absence of DNA (25). For DNA, we used a coarse-grained model in which three particles represent one nucleotide (26). Histone-DNA interactions include electrostatic interactions, excluded volume effects, and hydrogen bonds. These setups have been previously calibrated and already applied to investigate nucleosome dynamics (24). The nucleosome structure was modeled based on the crystal structures (PDB ID: 1KX5 (27) and 3LZ0 (28)) (Fig. 1B). We modeled the translocase as a torus-shaped excluded volume potential. The translocation was realized by applying a 14-pN force toward the upstream direction (half of the stall force [28 pN] of the bacterial RNA polymerase (29)) to the base-pair most proximal to the center of the torus. The force causes the next downstream base-pair to be pulled into the torus; when this base-pair becomes the closest to the torus center, the force is then applied to this new base-pair. The repetition of this procedure leads to a processive DNA translocation. Overall, the translocase (torus) relatively moves toward the nucleosome.

In the initial structure, the translocase was placed at 91 bps upstream from the dyad position of the nucleosome (Fig. 1C (i), Fig. 1D top left panel, and Fig. S2). As the simulation proceeded, the translocase collided with the nucleosome, unwrapping the nucleosomal DNA. Note that the position of the translocase (the center of the torus) was, on average, 7 bps behind that of the base-pair to be detached (Fig. 1D top left panel) due to the excluded volume of the translocase. When the translocase reached to −63 ± 2 bps and to −29 ± 5 bps, it stalled due to the relatively strong interaction between the base-pairs to be detached and the histone core complex (Fig. 1C (ii), (iii), Fig. 1D top left panel, and Fig. S2). We calculated the probability distribution of torus position from 80 simulation runs and found that the translocase statistically tended to stall at −57 ± 2 bps and −29 ± 5 bps before reaching the dyad (Fig. 1C top right panel), consistent with the previous reports (30–32).

Interestingly, after the translocase escaped from the −29 ± 5 bps stall, the DNA bps located at the dyad site (green curve in Fig. 1D top left panel) made a sudden and large transition, accompanied by the translocase movement (Fig. 1D top left panel). We repeatedly observed the repositioning in the 80 simulations (Fig. 1D bottom panels and Fig. S1). In these simulations, the dyad repositioning distance ranged from 78 to 102 bps (Fig. 1D and Fig. S1). In the repositioned nucleosome, the entry-side half of the histone core complex was wrapped by the right half of the Widom 601 sequence and the exit-side half by the random downstream DNA (Fig. 1A, Fig. 1C (iv), and Fig. S2). After the histone core complex repositioning, the translocase stalls at 20 ± 2 bps (−63 ± 2 bps from the new dyad position; Fig. 1C top left panel), supporting the almost complete nucleosome formation. In some trajectories, we observed a second repositioning event, though the limited length of the downstream DNA precludes the complete nucleosome formation (Fig. 1A and Fig. 1D bottom left panel).

### Relaxation simulation of partially unwrapped nucleosome

This sudden and long-distance (78 to 102 bps) repositioning cannot be explained by the pushing model in which one base-pair unwrapping is coupled to one base-pair histone core complex repositioning. In our simulations, the force was applied as soon as the nucleotide enters the torus. In reality, however, the force must be generated at a certain stage of the ATP hydrolysis cycle, which is much slower than our simulation timescale. Thus, the *in silico* translocation speed is much faster than reality in which the partially unwrapped nucleosome structure would relax every time the translocase moves one base pair. To study this process, we performed a second set of simulations starting from a partially unwrapped nucleosome structure obtained in the above simulations. In this case, instead of pulling, we fixed the base-pair at the center of the torus at its initial position.

First, we performed 80 simulation runs using the nucleosome conformation in which the translocase is at −18 bps as the initial structure (Fig. 2A top panel), which corresponds to just before the repositioning. As the simulation proceeds, the structure relaxed and significantly changed from the initial structure (Fig. 2A). This result suggests that indeed the structure did not relax enough in the pulling simulations. In the new set of simulations, we observed the downstream repositioning in 61% (49/80) of the simulation runs.

**Figure 2:**
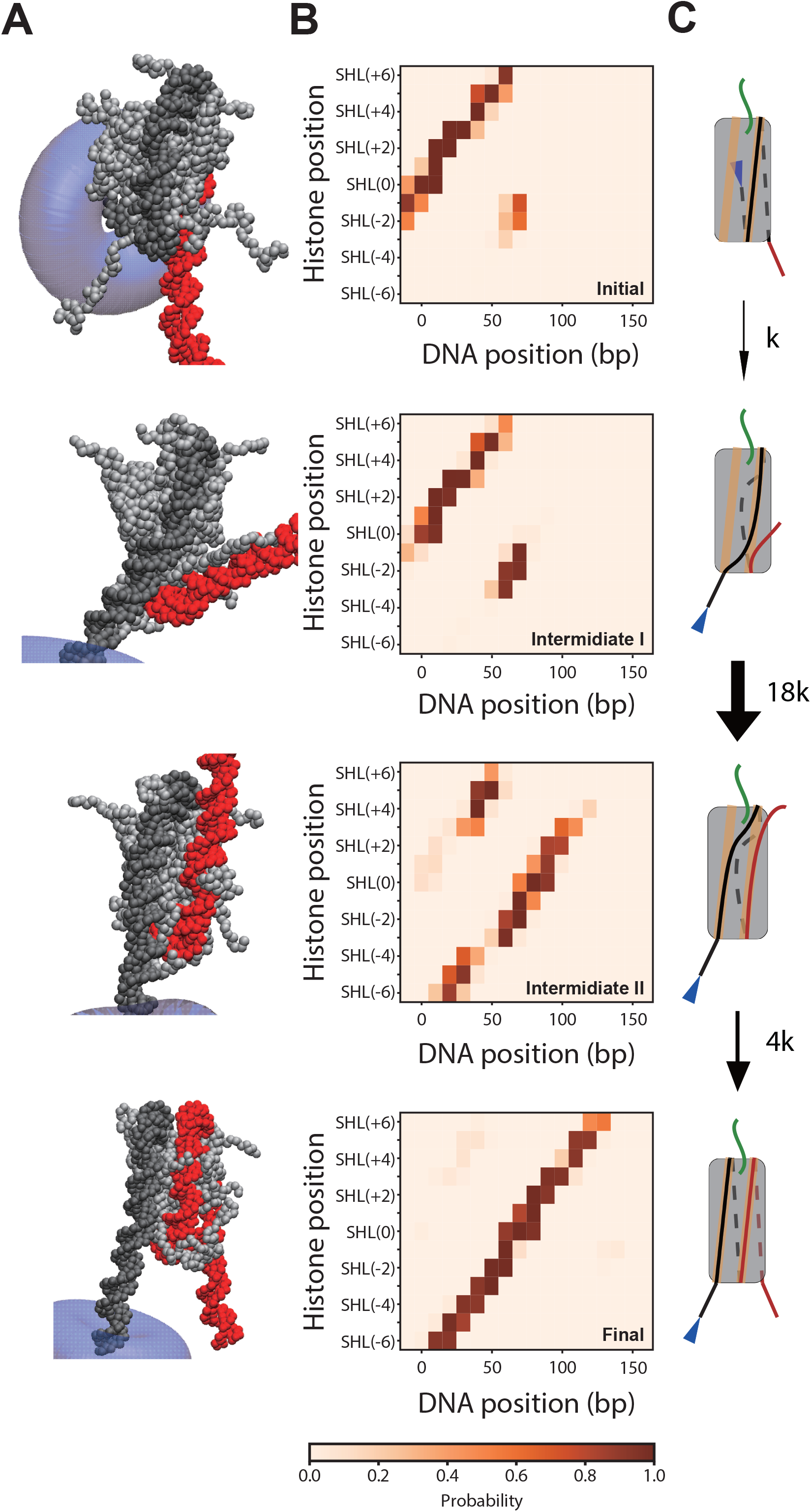
Relaxation simulations of the partially unwrapped nucleosome. (A) Representative structures in the simulation trajectories. DNA, the histones, and the translocase are colored according to the color scheme in Fig. 1C. (B) Contact maps showing contacts between a histone core complex and DNA. The maps were averaged over the cluster members. See Fig. 1B for the definition of super helical locations (SHL). (C) Cartoons explaining the lane-switch mechanism. The numbers next to the arrows represent relative transition rates.

Then, we sought to classify the structures in the 80 relaxation simulations. To achieve this, we calculated the contact maps of the histone core complex and DNA and performed a k-means clustering (Fig. 2B and Fig. S3C). The optimal numbers of clusters were decided to be seven based on the Calinski-Harabasz index (Fig. S4) (33), which is proportional to a ratio of inter-and intra-clusters dispersion. Visual inspection revealed that the seven clusters contain the initial conformation, an off-pathway intermediate, two on-pathway intermediates, and three final states. (Fig. 2B and Fig. S3). The main pathway contains four of the seven states: the initial, the two on-pathway intermediates I and II, and the final state (Fig. 2BC and Fig. S4A). In the initial state, −9th to 59th base-pairs contact with SHL (−1.5) to SHL (6.5) amino acids (Fig. 2BC top panel; See Fig. 1B for the positions of the super helical locations (SHL)). In the intermediate I state, the probability of contacts between −9th to 3rd base-pairs and SHL (−1.5) to SHL (0.5) amino acids decreases, and that between 59th to 71st base-pairs and the same amino acids increases (Fig. 2BC second panel). This result indicates the competitive binding of −9th to 3rd base-pairs and 59th to 71st base-pairs to the same amino acids. In the intermediate II state, the probability of contacts between 3rd to 34th base-pairs and SHL (0.5) to SHL (3.5) amino acids decreases, and that between the 19th to 40th base-pairs and SHL (−6.5) to SHL (−3.5) amino acids increases (Fig. 2BC third panel). In this state, the DNA originally in the exit side switched its lane (the binding site on the histone core complex). In the final state, the 18th to 145th base-pairs bind to SHL(−6.5) to SHL (6.5) amino acids, fully wrapping around the histone core complex (Fig. 2BC bottom panel). This state corresponds to the state after repositioning.

Based on the simulations, we computed the transition rates between the states along the repositioning pathway. Denoting the transition rate from the initial to the intermediate I states as *k* (=8.1×10^−9^ step^-1^), the rates from the intermediate I to the intermediate II states, and from the intermediate II to the final states are 19.7 ± 6.4 *k* and 4.0 ± 4.9 *k*, respectively (Fig. 2C and Fig. S2). This result suggests that spontaneous DNA dissociation after partial unwrapping is the bottleneck transition of downstream repositioning.

Next, we used the nucleosome structure in which the translocase is at −31 bps as an initial structure. In 93% (74/80) of these simulations, the histone core complex kept binding to the initial position (Fig. S5). In the remaining (6/80) simulations, H2A/H2B dimers dissociated from the H3/H4 tetramer. Interestingly, we did not observe downstream repositioning in these simulations. This is in sharp contrast to the simulations started at −18 bps, where we observed downstream repositioning in 61% (49/80) of the runs. Note that, when the translocase is at −31 bps and at −18 bps, the next base-pairs to be detached are at −19 bps and at −9 bps, respectively. Thus, the simulation results suggest that the detachment of −19th to −9th base-pairs from the histone core complex is essential for downstream repositioning.

Based on these simulations, we propose the lane-switch mechanism. In this mechanism, after a translocase unwraps nucleosomal DNA up to the site proximal to a dyad, the remaining wrapped DNA switches its binding region (lane) to that just vacated by the unwrapping, and then the downstream DNA rewraps, completing downstream repositioning. Next, we sought to validate this model experimentally.

### Restriction enzyme digestion assay

To validate the model, we chose T7 RNA polymerase (RNAP) as a model system. First, we designed 498 bps DNA substrates containing the T7 promoter and the nucleosome positioning sequence (Fig. 3A and Table S1). As the positioning sequence, the Widom 601 sequence was modified so that the non-template strand does not contain adenine from the entry nucleotide to one nucleotide before the stall site. The stall sites were chosen at −54 bps, −29 bps, and −14 bps (relative position from the center of the 601 sequence) (Fig. 3A and Table S1). MNase-seq assay (34) confirmed that this modification does not significantly affect nucleosome positioning (Fig. S6). When these DNA substrates were transcribed by the T7 RNA polymerase in the absence of adenine triphosphate in the buffer, transcription was terminated at these stall sites (Fig. S7A).

**Figure 3:**
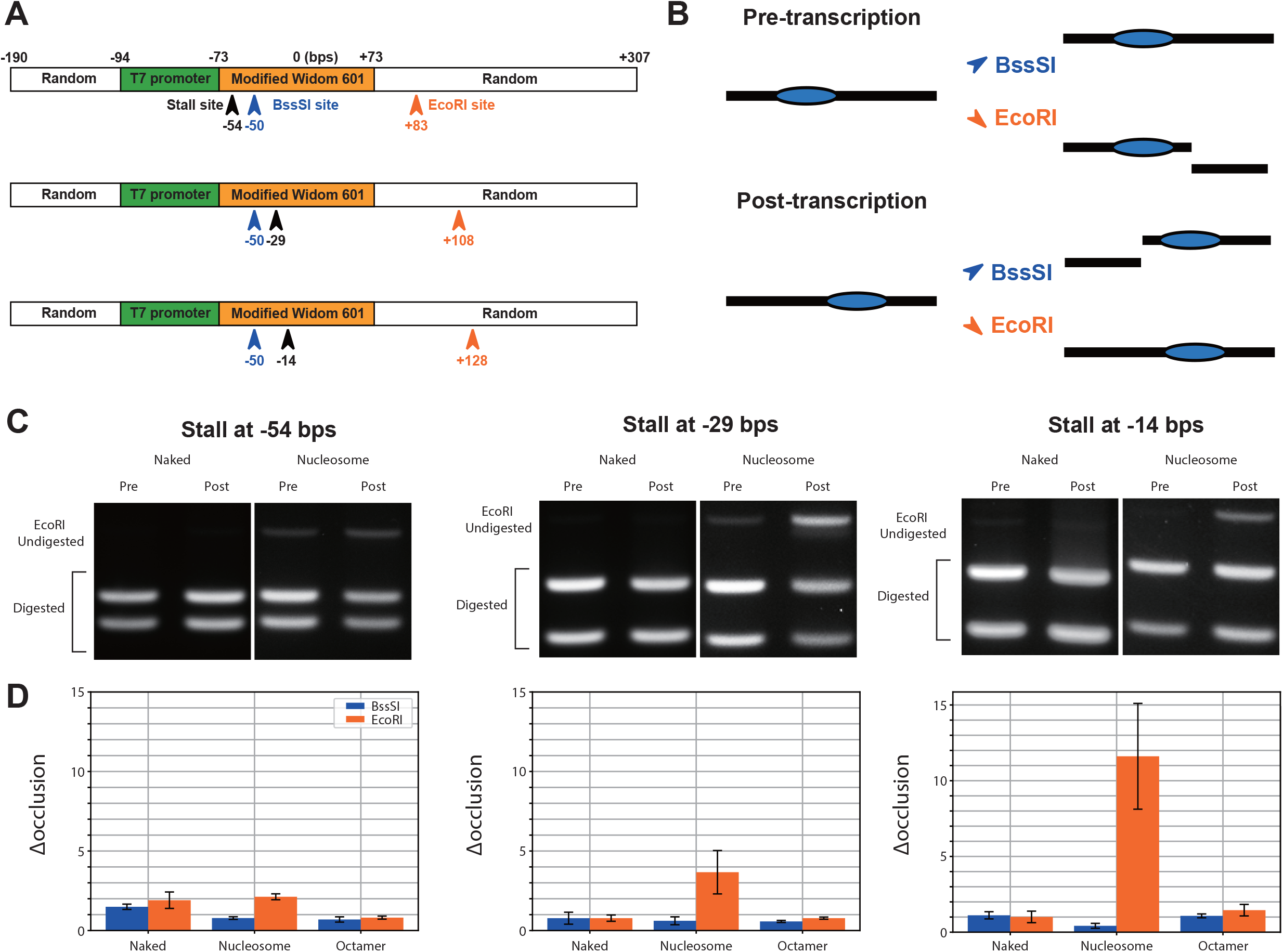
Restriction enzyme digestion assay. (A) The DNA sequence map of substrates used in the assay. The T7 RNAP stall sites where the translocase first encounter adenine are marked by the black arrows. BssSI and EcoRI restriction sites are marked by the blue and red arrows, respectively. (B) Cartoons of the experimental setup. At the pre-transcription stage, the BssSI site, but not the EcoRI site, is occluded by the histone core complex. On the other hand, if the histone core complex repositions upon transcription according to the lane-switch mechanism, the EcoRI site, but not the BssSI site, is occluded by the histone core complex. (C) Images of 1% agarose gel on which the EcoRI-digested products run. The DNA substrates were digested at the pre- and post-transcription stages. (D) Plot showing the intensity of the undigested band at pre-transcription stage divided by that of the post-transcription stage (Δ_*occlusion*_). The assay was repeated using naked DNA (Naked), nucleosome reconstituted DNA (Nucleosome), and the naked DNA with histone core complexes in solution (Octamer).

The designed sequences contain the BssSI restriction site in the modified 601 sequence, which is supposed to be occluded by a histone core complex (35). The sequences also contain the EcoRI restriction, which is supposed to be occluded when the histone core complex repositions downstream upon transcription. Thus, the repositioning frequency can be quantified by the restriction enzyme digestion assay (Fig. 3B). In this assay, we digested the DNA substrates with BssSI and EcoRI and run the products on 1% agarose gel (Fig. S8). The digested and undigested fragments were separated on a gel, and the intensities of the undigested product were measured (Fig. 3C). We defined the Δ_*occlusion*_ as the intensity after transcription divided by the intensity before transcription.

For the naked DNA, the values of Δ_*occlusion*_ were about one irrespective of the enzymes and the stall sites (Fig. 3D and Fig. S8), suggesting that the enzymes can bind to the restriction sites when the RNAP stalls at −54 bps, −29 bps, or −14 bps. When a nucleosome was reconstituted on the DNA substrates, Δ_*occlusion*_ of EcoRI after transcription was 2.1 ± 0.2, 3.7 ± 1.4, and 11.6 ± 3.5 for stalling at −54 bps, −29 bps, and −14 bps, respectively (Fig. 3D). The micrococcal nuclease (MNase) assay confirmed nucleosome formations before and after transcription (Fig. S7B). These results suggests that the histone core complex repositions downstream when RNAP proceeds to −14 bps. On the other hand, Δ_*occlusion*_ of BssSI was 0.8 ± 0.1, 0.6 ± 0.2, and 0.4 ± 0.2 for stalling at −54 bps, −29 bps, and −14 bps (Fig. 3D). Thus, the BssSI site in the nucleosome positioning sequence was more exposed when RNAP proceeds to −14 bps, as expected from an increased probability of downstream repositioning.

To check if a nucleosome repositions by temporal dissociation and re-association of a histone core complex, we performed the same assay using naked DNA in the presence of histone core complexes in solution. As a result, Δ_*occlusion*_ was about one irrespective of the enzymes and the stall sites (Fig. 3D), suggesting that, once in solution, the histone core complexes do not occlude the BssSI or EcoRI sites. This result argues against the temporal dissociation and re-association as the mechanism for downstream repositioning.

Together, the enzyme digestion assays supports our model of histone core downstream repositioning coupled with DNA unwrapping.

### Deep sequencing assay

In the restriction enzyme digestion assay, we could not identify the position of the histone core complex down to base-pair resolution. To localize the complex, the substrates before and after the transcription was digested by MNases, and the digested products were sequenced using a nanopore sequencer (36). MNases do not efficiently digest the DNA wrapping around a histone core complex. Thus, we can use the DNA reads from the nanopore sequencer as a proxy for a nucleosome position.

First, a nucleosome was reconstituted using the DNA substrate described above (Fig. 3A). Then, the substrate was digested, purified, and sequenced. The distribution of the read length shows a single peak around 158 bps (Fig. S9). This result indicates that the sequenced products wrap around histone core complexes before the purification. We define the position of the central base-pair of the read as a nucleosome position. Only DNA reads with length from 150 to 166 base pairs are considered for the analysis.

Next, after the nucleosome reconstitution, the T7 RNAP transcriptions were induced in the absence of adenine in solution. The DNA substrates contain the stall site at −14 bps or −54 bps as described above. After transcription, the substrates were digested, purified, and sequenced.

In the case of the stall position at −54 bps, the frequency distribution of the nucleosome position shows a peak around 50 bps (Fig. 4A). This peak bears two interpretations: i) unwrapping of each base pair is coupled with one base pair repositioning of the histone core complex (pushing), and ii) the histone core complex spontaneously slides to this meta-stable positioning site due to the occlusion of the modified 601 sequence by the T7 RNAP (spontaneous sliding). However, when RNAP proceeds up to −54 bps, the pushing model would predict repositioning by up to 19 bps, which is inconsistent with the 50 bps peak (Fig. 4E).

**Figure 4:**
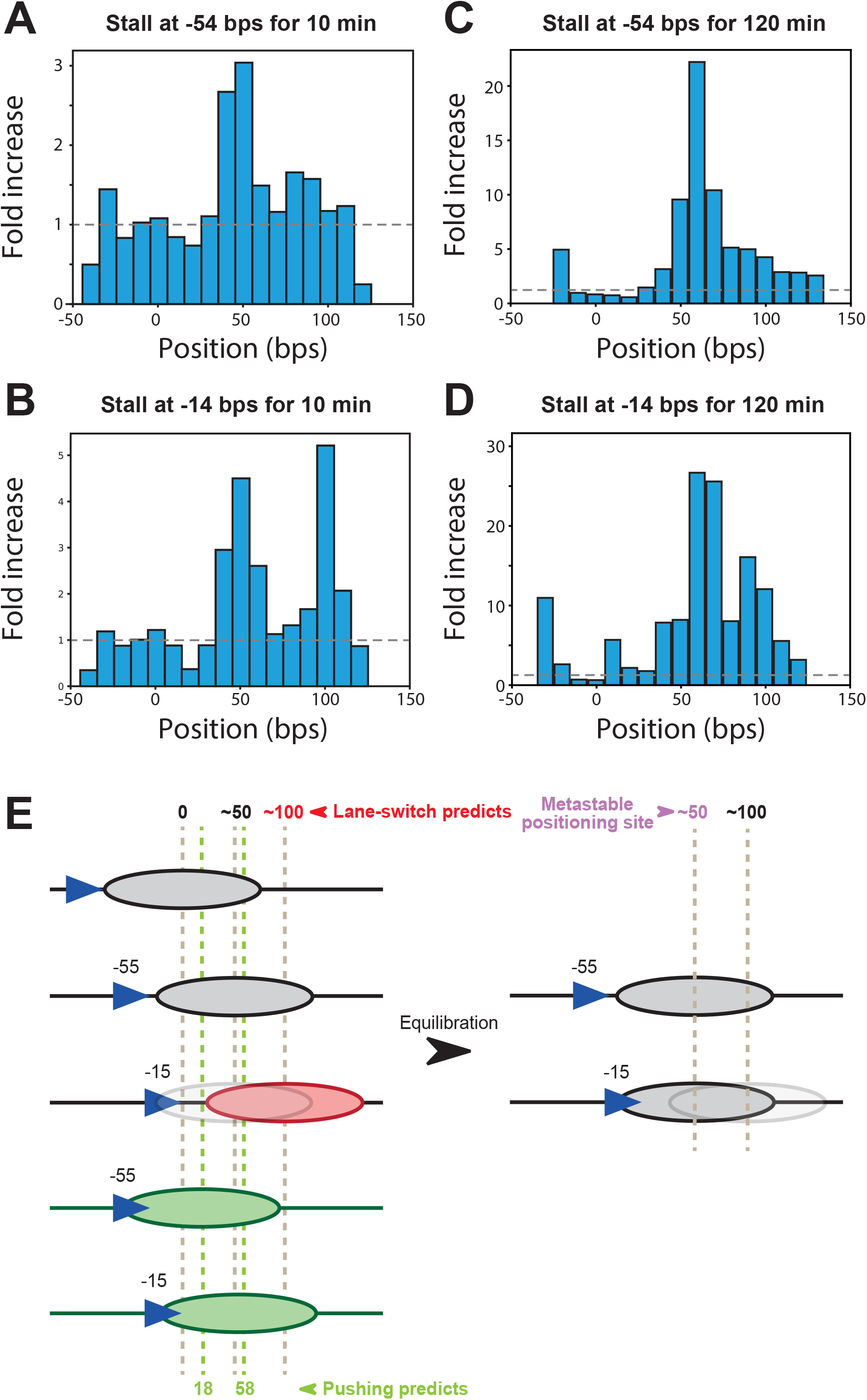
MNase-seq assay. (A to D) Frequency distribution of a nucleosome position at the post-transcription stage. The origin corresponds to the position of the center base-pair of the Widom 601 sequence. The distribution was normalized by dividing by the distribution at the pre-transcription stage. The RNAP stalls at −54 bps for 10 min (A), at −14 bps for 10 min (B), at −54 bps for 120 min (C), and at −14 bps for 120 min (D). (E) Cartoons of interpretation of the experimental data.

Interestingly, in the case of the stall position at −14 bps, the frequency distribution of the reposition distance shows two peaks (Fig. 4B) around 50 bps and 100 bps. When RNAP proceeds up to −14 bps, the pushing model predicts repositioning by 59 bps, which is inconsistent with the 100 bps peak. On the other hand, the 100 bps peak (101 ± 8 bps in Fig. 4B) is within the 78-102 bps range of the repositioning distance predicted by the lane-switch mechanism (Fig. 4E). Thus, we concluded that the 100 bps peak in the frequency distribution of reposition distance supports the lane-switch mechanism.

Next, we consider the mechanism behind the 50 bps peak, which was observed irrespective of the stall position. This peak is best explained by the spontaneous sliding model because of the following reasons: (i) If the unwrapping of each base pair were coupled with the one base pair reposition of the histone core complex, the frequency distribution should show peaks around 19 bps and 59 bps in the case of stalling at −54 bps and −14 bps, respectively, which were not observed. (ii)The theoretically predicted free energy landscape of the nucleosome positioning shows a meta-stable basin around 50 bps (Fig. S10), indicating that the site can become the preferred destination site for the spontaneous sliding once the RNAP destabilizes the original 601 positioning at 0 bp. (iii) When RNAP stalling at −15 bps is prolonged (for 120 min instead of 10 min), giving the histone core more time to slide toward its equilibrium position, the relative heights of the two peaks changes, increasing for the 50 bps peak and decreasing for the 100 bps peak (Fig. 4B and Fig. 4D). Also, when RNAP stalls at −54 bps for 120 min, the 50-bps peak gets sharper (Fig. 4A and Fig. 4C), supporting that this is one of the stable destination sites for the spontaneous sliding. These considerations make us conclude that the 50 bps peak can be explained by the spontaneous sliding model. Despite the existence of such metastable sites, RNAP repositions the histone core complex to the distal 100 bps sites when it unwraps up to −15 bps but not −55 bps (Fig. 4E), supporting the lane-switch mechanism.

Together, the MD simulations, the enzyme digestion assays, and the MNase-seq assays consistently suggest that the histone core complex repositions downstream through the lane-switch mechanism.

## DISCUSSION

In this study, we sought to elucidate the molecular mechanism by which the histone core complex repositions when a translocase unwraps nucleosomal DNA. To achieve this, we performed MD simulations, restriction enzyme digestion assays, and deep sequencing assays. Our MD simulations in which a model translocase unwraps nucleosomal DNA (Fig. 1) revealed that the histone core complex repositions downstream upon unwrapping. Further structural relaxation simulations (Fig. 2) revealed the detailed molecular mechanism of downstream repositioning. In this mechanism, after a translocase unwraps nucleosomal DNA up to the site proximal to the dyad, the remaining wrapped DNA switches its binding region (lane) to that vacated by the unwrapping, and finally the downstream DNA rewraps, completing downstream repositioning. The enzyme digestion assays (Fig. 3) and the deep sequencing assay (Fig. 4) experimentally supported the mechanism.

Previous studies proposed the Pol II-type (9–16) and Pol III-type (9, 17–21) molecular mechanisms of histone core complex repositioning upon nucleosome unwrapping. In the Pol II-type mechanism, the translocase passes through the nucleosome without changing the position of the histone core complex. To realize this, the histone core complex electrostatically interacts with the translocase, forming the Ø-intermediate structure. On the other hand, in the Pol III-type mechanism, a histone core complex repositions upstream. Notably, the Pol III-type mechanism was proposed based on experiments in which the nucleosome was reconstituted at the downstream end of DNA, precluding downstream repositioning (9, 17–21). In the current work, we removed this constraint by reconstituting nucleosome at the middle of the DNA substrate, showing that the histone core complex repositions downstream upon nucleosome partial unwrapping. Whether the histone core complex repositions downstream or upstream may depend on the availability of DNA, which is regulated by its persistence length and occlusion by other proteins.

Gottesfeld and Luger reported that T7 RNAP stalls at the entry base-pair when the pyrrole-imidazole polyamides binding to the dyad, preventing the histone core complex repositioning (22). This result supports the mechanism by which one base-pair unwrapping is coupled to one base-pair repositioning (the pushing model). However, the study did not fully exclude the possibility that the polyamide spatially proximal to the entry site directly affects the RNAP translocation. In fact, it was reported that T7 RNAP can partially unwrap nucleosomal DNA without histone core complex repositioning (9, 17–20). Also, the current study shows that repositioning does not take place until a translocase partially unwraps nucleosomal DNA. Though the current study does not fully exclude the possibility that the repositioning based on the pushing mechanism take place, in our deep sequencing assay, we clearly observed repositioning which cannot be explained by the pushing mechanism but can be explained by the lane-switch mechanism.

In the current study, we proposed that the nucleosome repositioning according to the lane-switch mechanism when there is little attractive interaction between a translocase and a histone core complex. This indicates that other types of repositioning mechanisms should be based on an intricate interaction network between the translocase and the histone core complex. For example, Pol II can bypass a nucleosome without histone core complex repositioning because of strong electrostatic interaction between the translocase and the complex (15). The lane-switch mechanism represents a fundamental passive mechanism useful for understanding the molecular basis of transcription through nucleosome, histone recycling, and nucleosome remodeling, where more intricate interaction between translocase and the histone core complex must be involved in.

## METHODS

### Coarse-grained MD Simulations

We used the AICG2+ model (37) for a histone octamer, and the 3SPN.2C model (26) for DNA. In the AICG2+ model, each amino acid is represented by one bead placed on the C_α_ atom of each residue. In the 3SPN.2C model, each nucleotide is represented by three beads placed at the centers of mass of phosphate, sugar, and base groups. The nucleosome structure was modeled based on the crystal structures (PDB ID: 1KX5 (27) and 3LZ0 (28)) (Fig. 1B), and 32 and 96 bps DNA strands were added upstream and downstream of the nucleosome, respectively (Table S1). The same potential energy functions as previous studies were used to stabilize the nucleosome structure (24).

We modeled a translocase as a torus. The potential energy function of the excluded volume term of the torus (Fig. S9) is

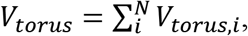

where

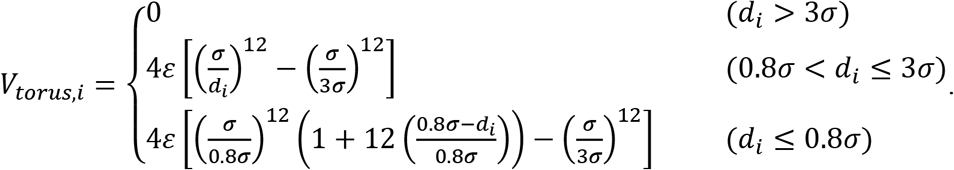

The distance between the surface of a torus and a particle in cylindrical coordinates is

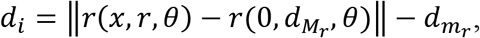

where 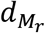 is the major radius of the torus, and 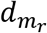 is the minor radius of the torus. In all simulations, we set the potential parameters *σ, ε*, 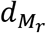, and 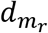 to 1.0 Å, 1.0 kcal/mol, 35 Å, and 22.5 Å, respectively.

The torus through which the upstream DNA passes was fixed in space. In the initial structure, the −19th base-pair is at the center of the torus. Translocation was realized by applying a 14-pN force in the upstream direction (half of the stall force [28 pN] of the bacterial RNA polymerase (29)) to the base-pair closest to the center of the torus (six DNA beads; 2.3 pN per each). This procedure leads to a processive motion of the torus-like translocase along DNA toward the nucleosome.

We performed the 80 pulling simulations and the 80 relaxation simulations without pulling for each stall position (−54 bps or −14 bps) using the software CafeMol 3.2 (https://www.cafemol.org) (38). The equations of motion were integrated over time by Langevin dynamics with the time step set to 0.3 cafemol time (∼14.7 fs). The mono-valent ion concentration in the Debye-Huckel model was set to 300 mM.

### Restriction enzyme digestion assay

The T7_601_stall_-54, T7_601_stall_-29, and T7_601_stall_-14 DNA substrates were chemically synthesized (Eurofins genomics; Table S1) and amplified by polymerase chain reactions. We purified budding yeast histone proteins (H2A, H2B, H3, and H4) using an E. coli protein expression system and reconstituted nucleosomes using the purified proteins and the amplified DNA substrates as described previously (39). The reconstitusion was confirmed by MNase assays in which MNases (M0247S, New England BioLabs) digest linker DNA for 30 minutes and the digested products were run on 1% agarose gel. In vitro transcription on the DNA substrates was performed by using HiScribe T7 High Yield RNA synthesis kit (E2040S, New England BioLabs) for 120 min at 37 °C in the presence of 1 mM GTP, 1 mM CTP, and 1 mM UTP when otherwise specified. Next, we added 1 μL of RNase I (M0243, New England BioLabs) to reactions in 1 x NEBuffer 3.1 (B7003, New England BioLabs) and incubated for 30 minutes at 37°C to remove the transcripts. Then, we exchanged the buffer for the CutSmart buffer (B7204 New England BioLabs) using a 10K MWCO Amicon® Ultra-0.5 Centrifugal Filter (UFC510024, Merck), added BssSI-v2 (R0680, New England BioLabs) or EcoRI-HF (R3101, New England BioLabs), and incubated for 15 minutes at 37 °C. The digested products were extracted using a phenol:chloroform:isoamyl (25:24:1) solution (Nakalai Tesque) and run on 1% agarose gel. The gel was imaged using the iBright FL1500 imaging system (Thermo Fisher Scientific). The images were analyzed using ImageJ software (40).

### Deep sequencing assay

The nucleosome reconstitution, transcription, and MNase digestion procedures are the same as described above. After the MNase digestion, we purified the digested products using a spin column (A9281, Promega) to avoid organic solvent contamination. We prepared a DNA library using the ligation kit (SQK-LSK109, Oxford Nanopore) and loaded the library into the MinION R9.4.1 nanopore flowcell (Oxford Nanopore) according to the manufacturer’s instruction. The signals from the device were analysed using MinKNOW software (Oxford Nanopore) to obtain sequence reads.

### Free energy of nucleosome assembly

The free energy of nucleosome assembly along the DNA sequence was estimated using a Markov model described in Ref. (41), which is based on Monte-Carlo simulations of nucleosomes (42) at 300 K using a rigid-base pair model of DNA (43). To estimate the free energy of nucleosome assembly, we only take into account the DNA section up to the last contacts with the central H3/H4 tetramer (+/-28 bps away from the dyad), since both our tests and other studies found this to be a better predictor of nucleosome positioning along DNA (44).

## AUTHOR CONTRIBUTIONS

T.T. designed the study. F.N. prepared all the materials. F.N. and S.T. designed the molecular dynamics simulations. F.N. performed molecular dynamics simulation, restriction enzyme digestion assay, MNase-seq assay. All authors discussed the findings and co-wrote the manuscript.

## ACKNOWLEDGMENTS

We thank members of the theoretical biophysics laboratory at Kyoto University for discussions and assistance throughout this work. Also, we thank Prof. Hitoshi Kurumizaka and his laboratory members for helping us to purify histones and reconstituting nucleosome. This work was supported by PRESTO (JPMJPR19K3; to T.T.), Grant-in-Aid for Scientific Research (B; 18H06046; to T.T.), Grant-in-Aid for Scientific Research on Innovative Areas (Molecular engine; 19H05392; to T.T.), Grant-in-Aid for Scientific Research on Innovative Areas (Chromatin potential; 19H05260; to T.T.), and the CREST grant of Japan Science and Technology Agency (JST) (JPMJCR1762; to S.T.).

## FIGURE LEGENDS

**Figure S1:**
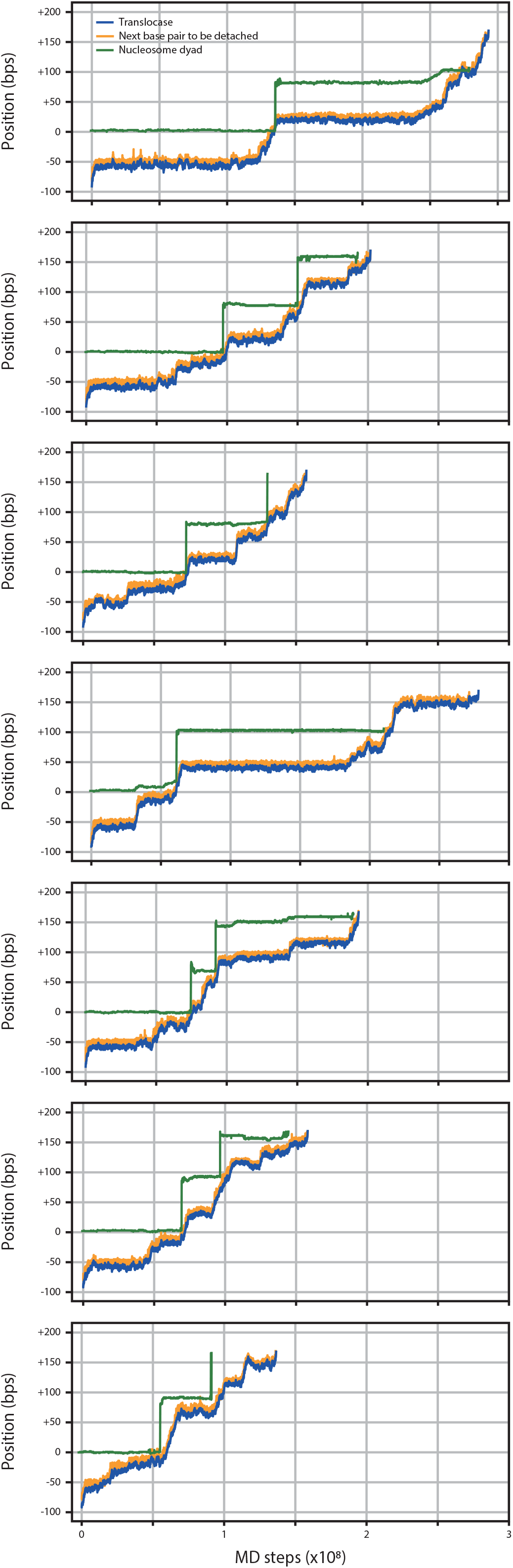
Representative simulation trajectories of the positions of the translocase, the base-pair to be detached, and the dyad.

**Figure S2:**
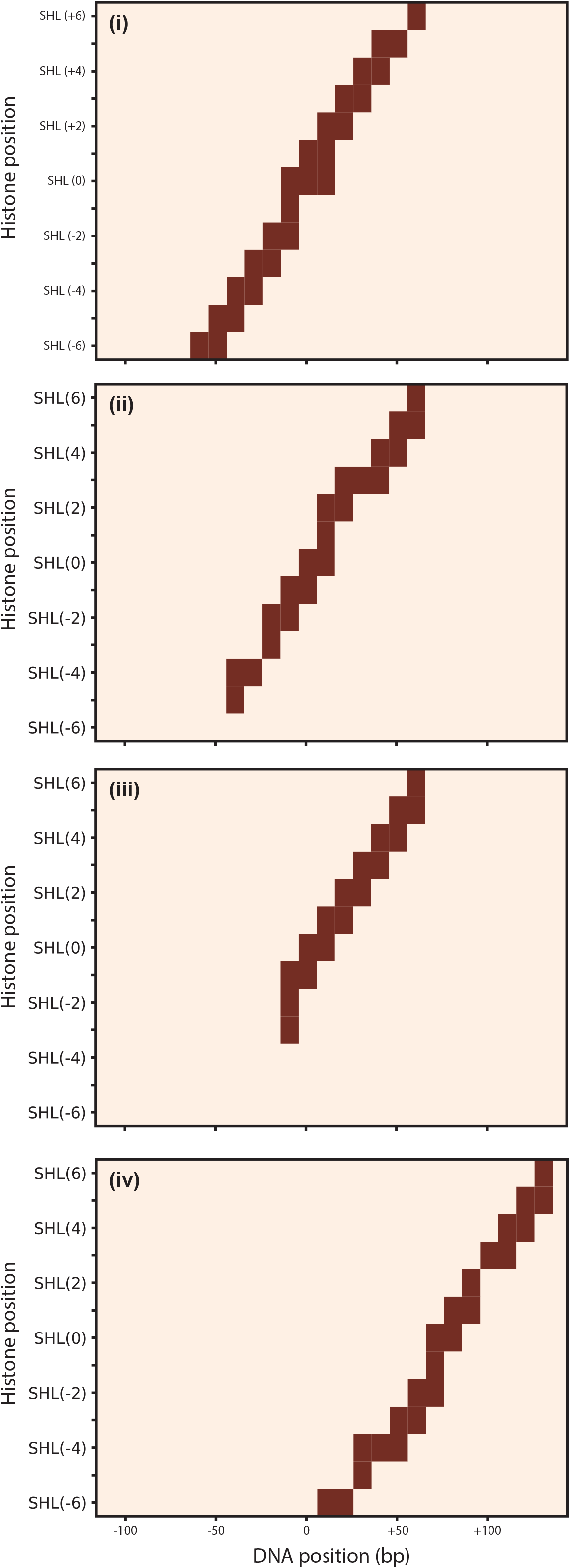
Contact maps showing contacts between a histone core complex and DNA. The maps were calculated using the structures from the simulation trajectory in Fig. 1D top left panel. See Fig. 1B for the definition of super helical locations (SHL).

**Figure S3:**
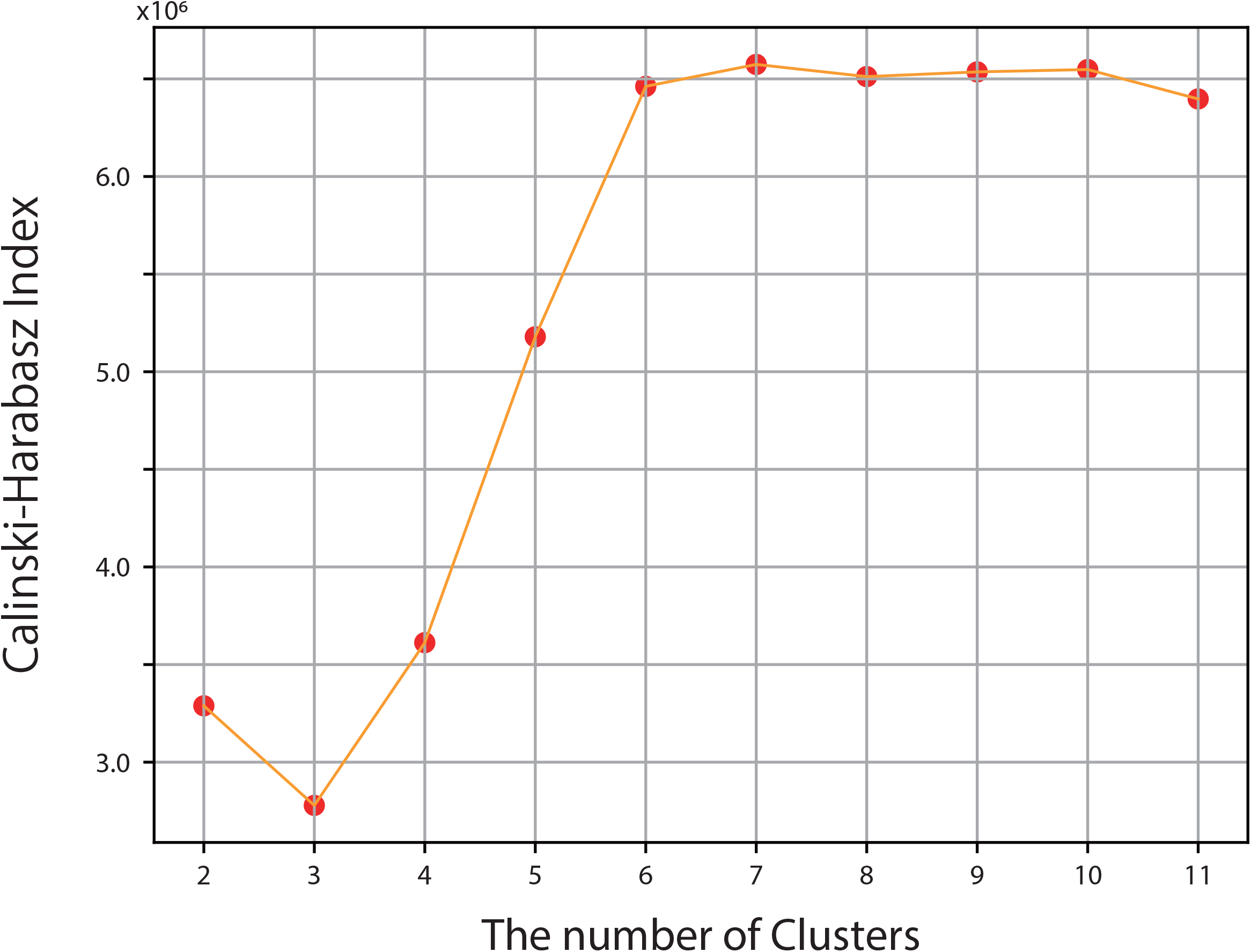
Calinski-Harabasz indexes depending on the number of clusters. Clustering of the contact maps from the relaxation simulation was performed by the k-means algorithm.

**Figure S4:**
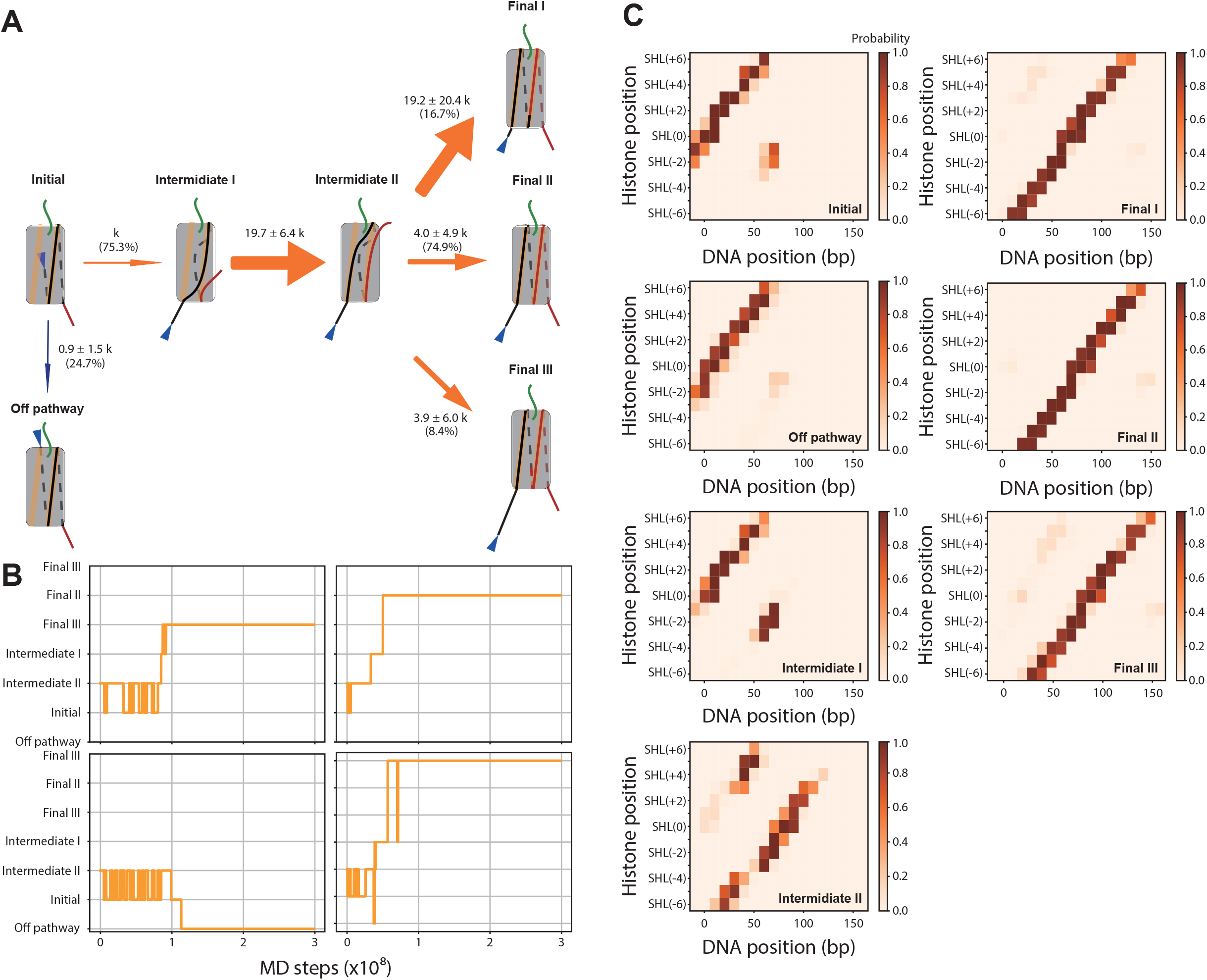
Relaxation simulations of the partially unwrapped nucleosome. (A) Schematic diagram of state transitions in the simulations. Numbers near arrows and those in parentheses represent transition rates and branching ratios, respectively. The color scheme is the same as in Fig. 2C. (B) Representative time trajectories of state transitions. The transition rates in (A) were calculated using the trajectories. (C) Contact maps of all the clusters from relaxation simulations showing contacts between histone core complex and DNA. The maps were averaged over the cluster members.

**Figure S5:**
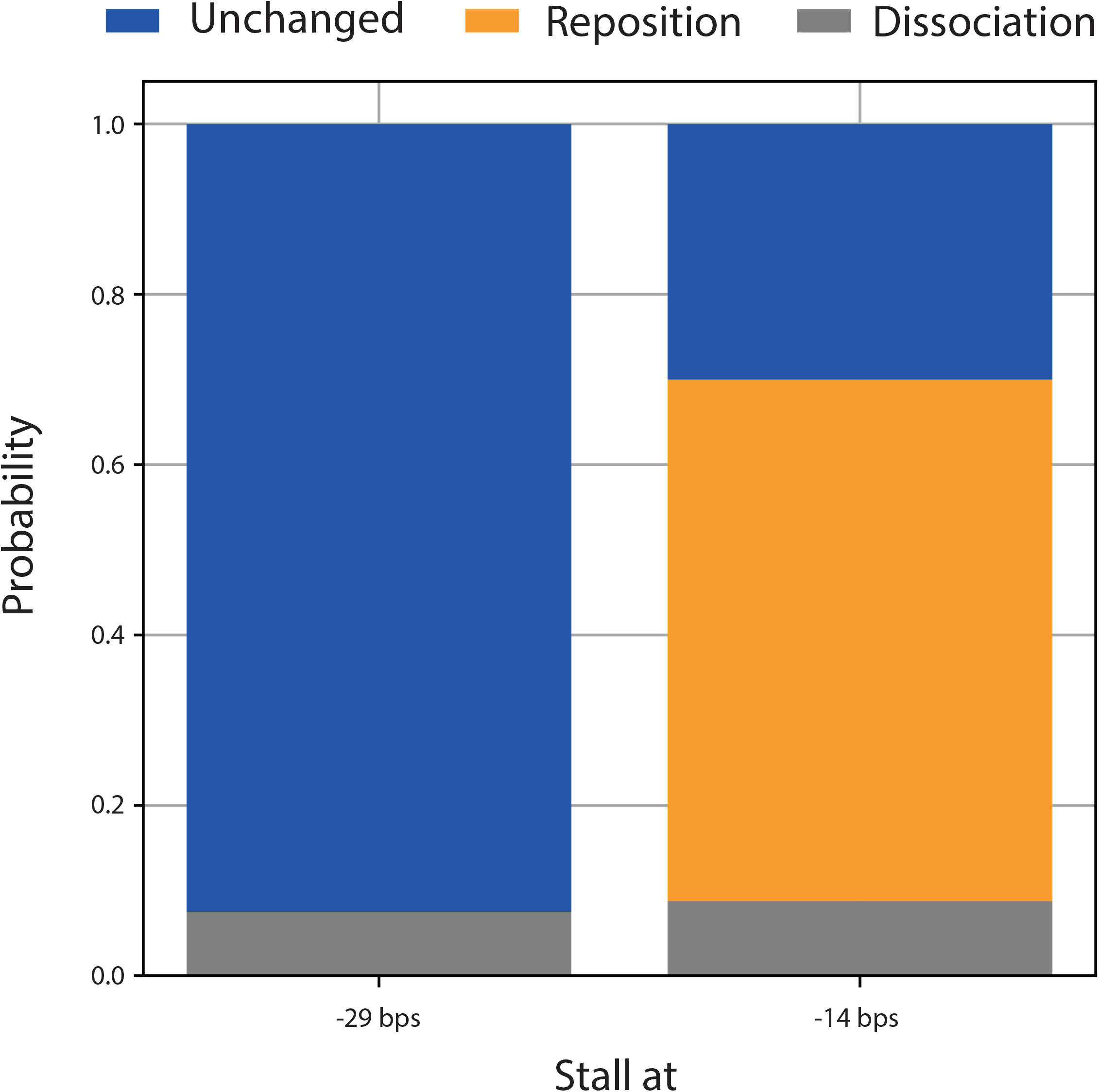
Probabilities of nucleosome states after 3.0 × 10^8^ steps of relaxation simulations when translocase stalls at −31 and −18 bps.

**Figure S6:**
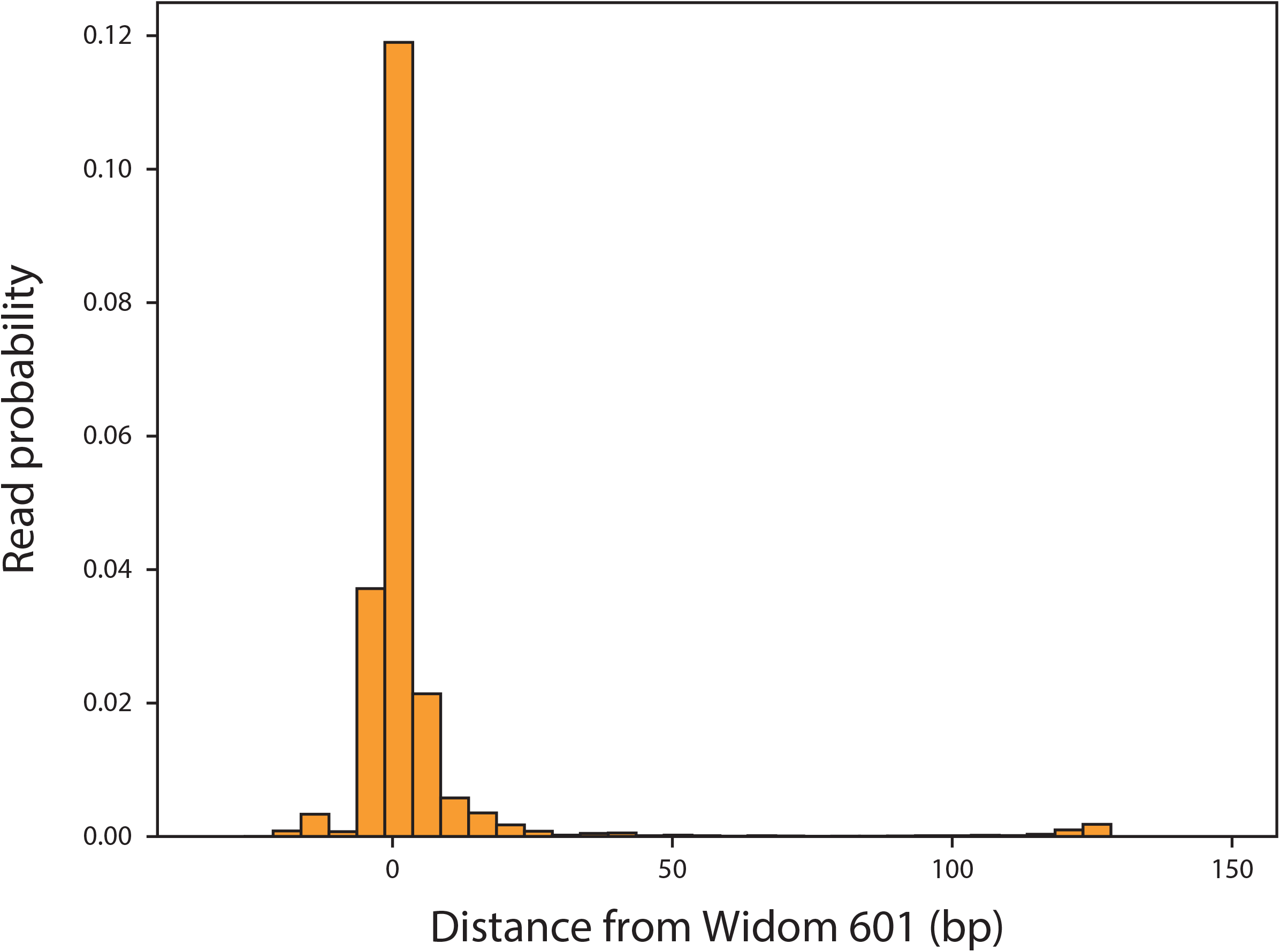
Probability distribution of the read position. The DNA substrate containing the modified Widom 601 sequence reconstituted into nucleosome was digested and sequenced (MNase-seq). The origin corresponds to the center base pair of the 601 sequences.

**Figure S7:**
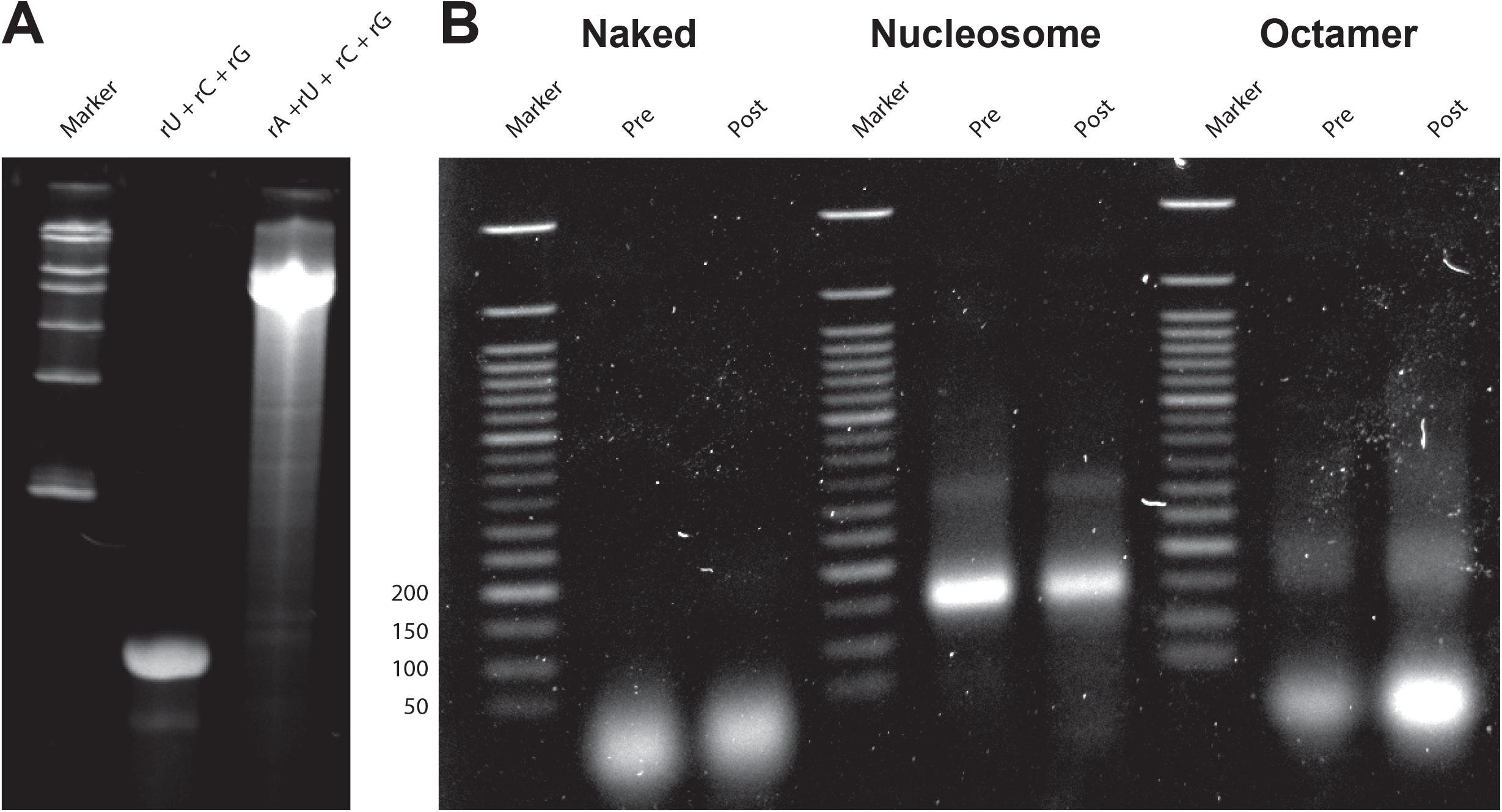
(A) Transcription RNA products run on 15% urea polyacrylamide gel. The transcription assays were repeated in the presence of UTP, CTP, GTP (rU + rC + rG) and in the presence of ATP, UTP, CTP, and GTP (rA + rU + rC + rG). The marker is RiboRuler Low Range RNA Ladder (SM1831, Thermo Scientific). (B) MNase assay of the transcribed substrates. The DNA substrates at the pre-and post-transcription stages were digested by MNase for 30 min and run on 1% agarose gel. The assay was repeated using naked DNA (Naked), nucleosome reconstituted DNA (Nucleosome), and the naked DNA with histone core complexes in solution (Octamer). The marker is Quick-Load Purple 50 bp DNA Ladder (N0556S, New England BioLabs).

**Figure S8:**
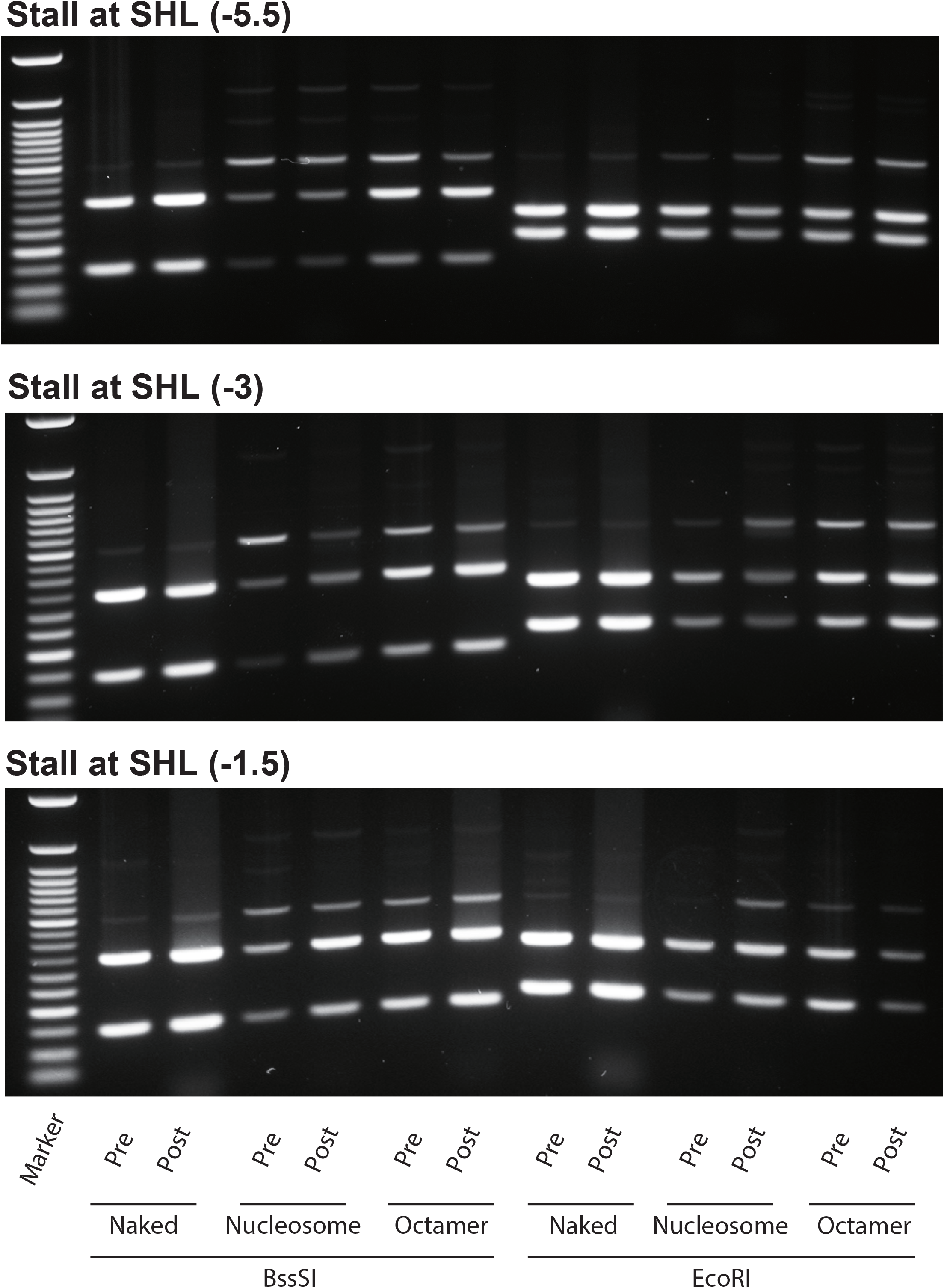
The images of 1% agarose gel on which the digested products run. The DNA substrates were digested by BssSI and EcoRI at the pre-and post-transcription stages. Depending on the substrates, the RNAP stalled at −54 bps (top), −29 bps (middle), and −14 bps (bottom), respectively. The assay was repeated using naked DNA (Naked), nucleosome reconstituted DNA (Nucleosome), and the naked DNA with histone core complexes in solution (Octamer). The marker is Quick-Load Purple 50 bp DNA Ladder (New England BioLabs; N0556S).

**Figure S9:**
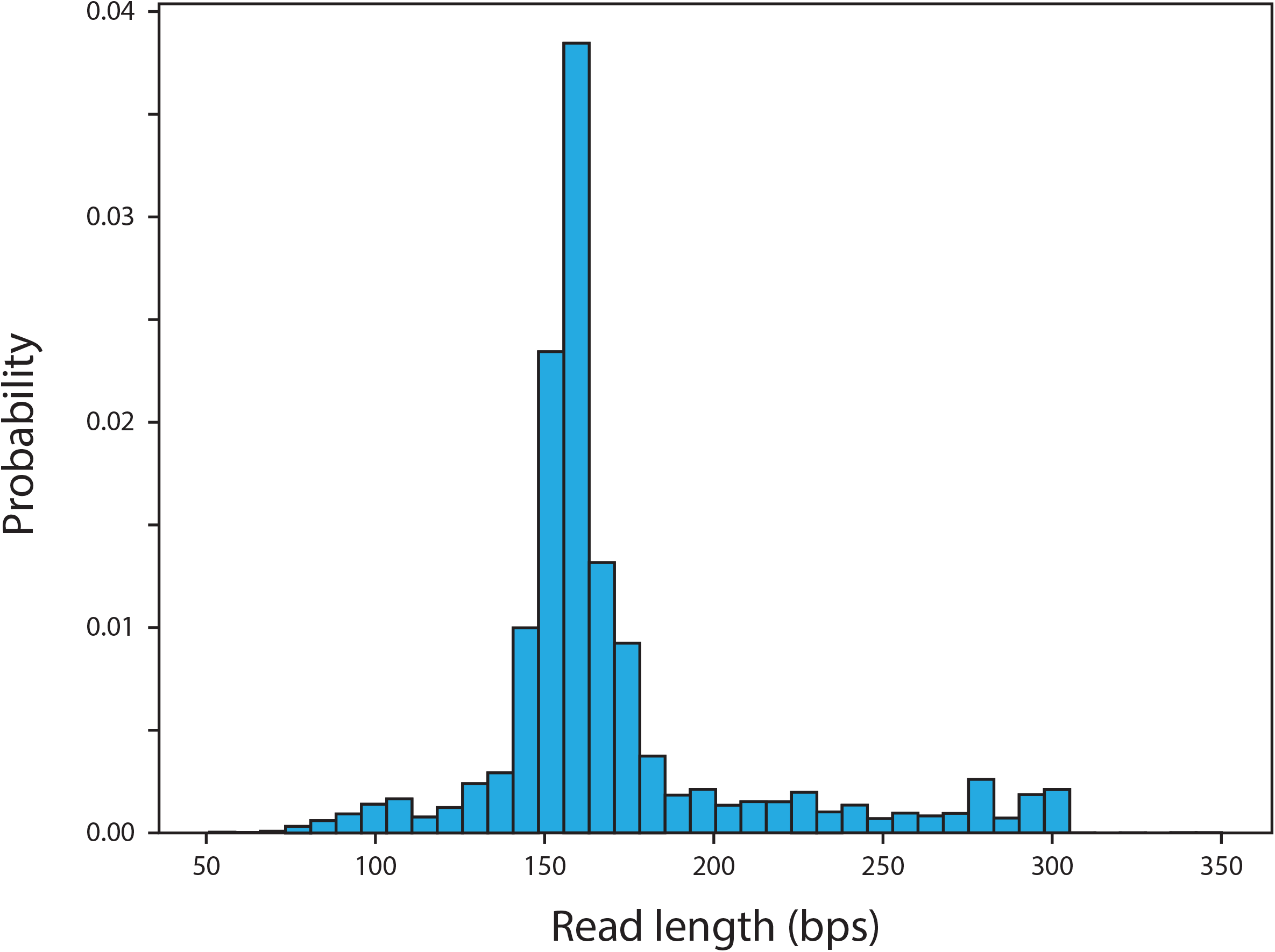
Probability distribution of the read length of MNase-seq assay. The DNA substrate containing the modified Widom 601 sequence reconstituted into nucleosome was digested and sequenced.

**Figure S10:**
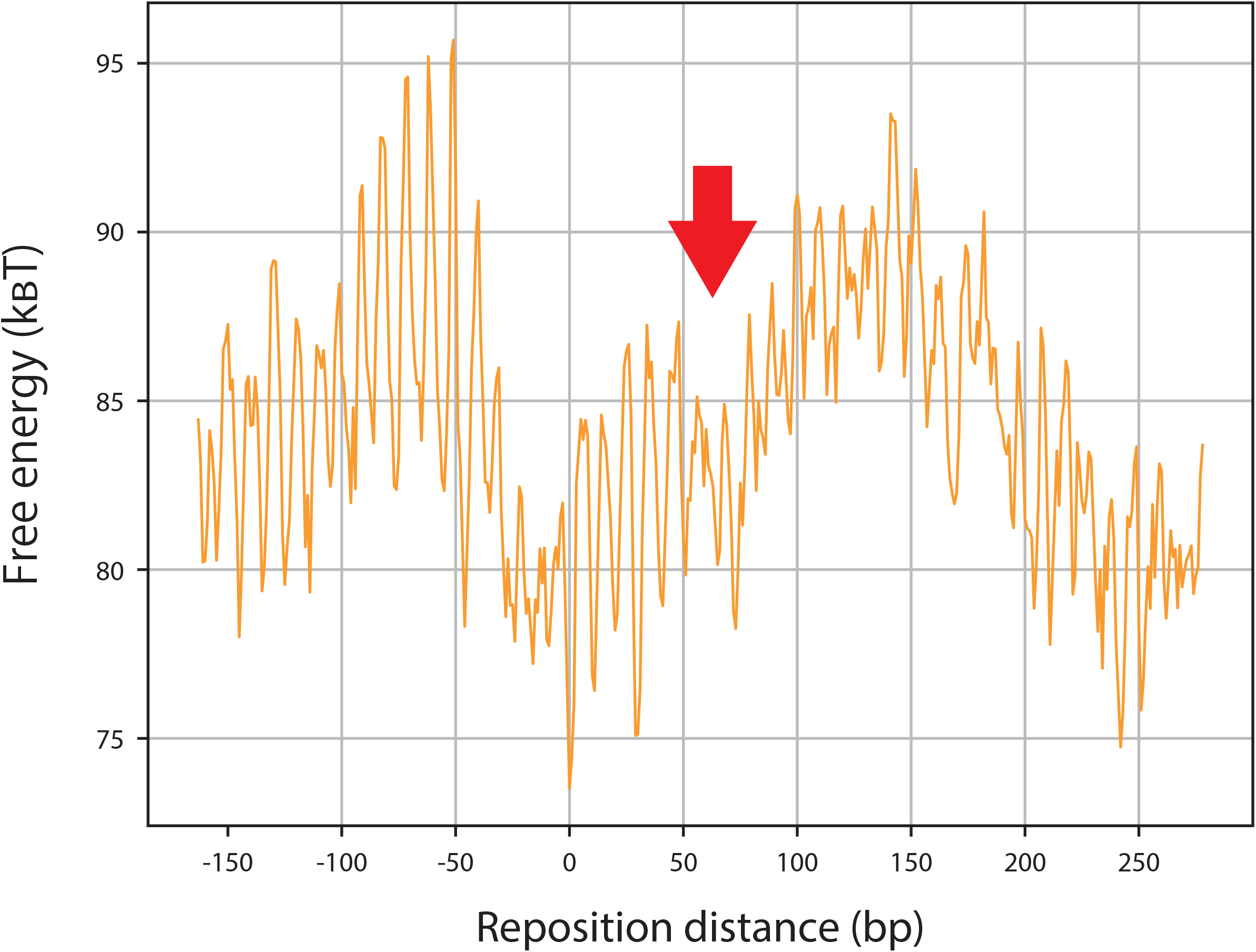
The predicted free energy landscape of a nucleosome repositioning. The red arrow points to the position of the meta-stable region.

**Figure S11:**
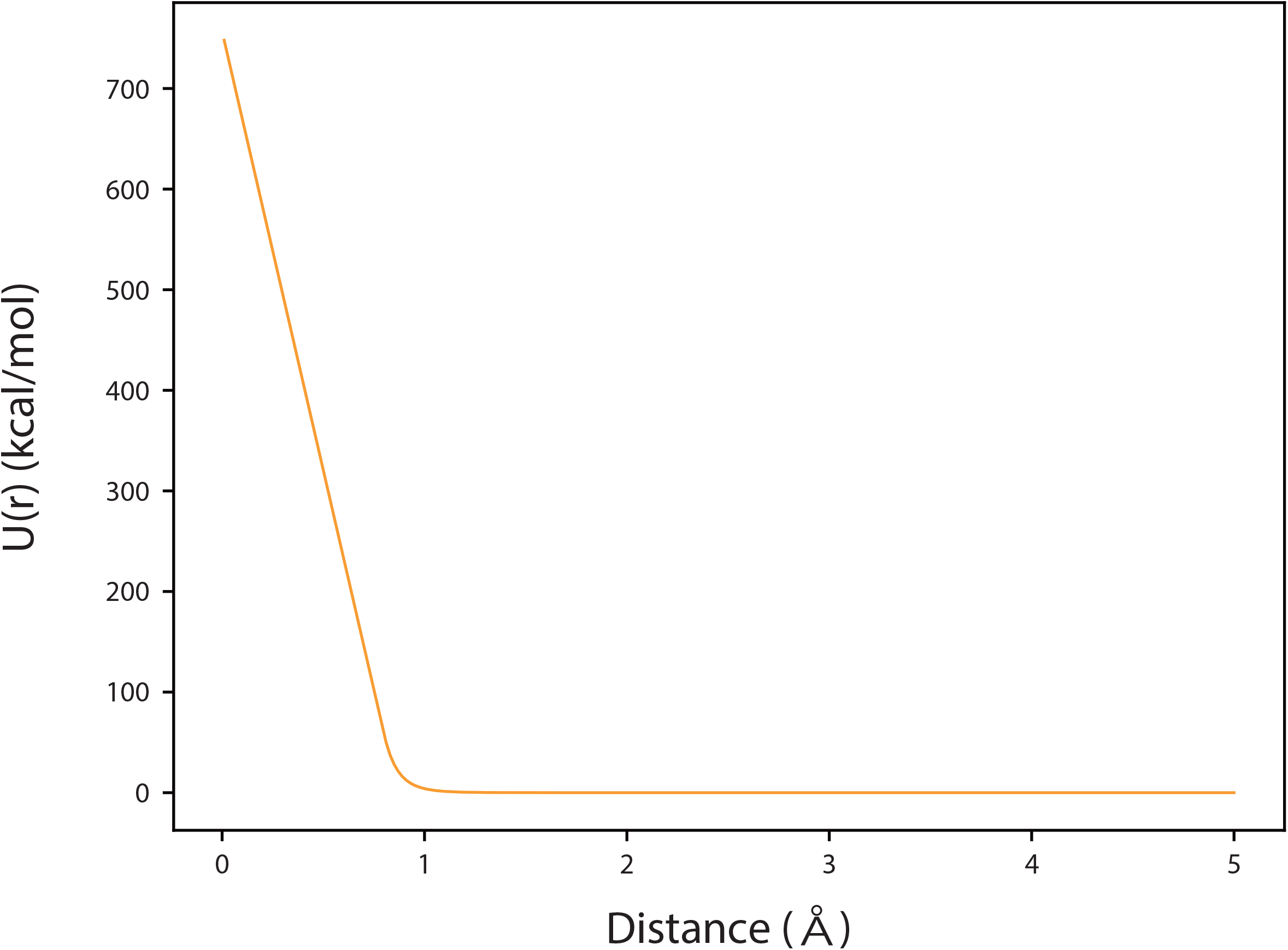
The potential energy function describing excluded volume effect of a translocase modeled as a torus.

**Table S1:**
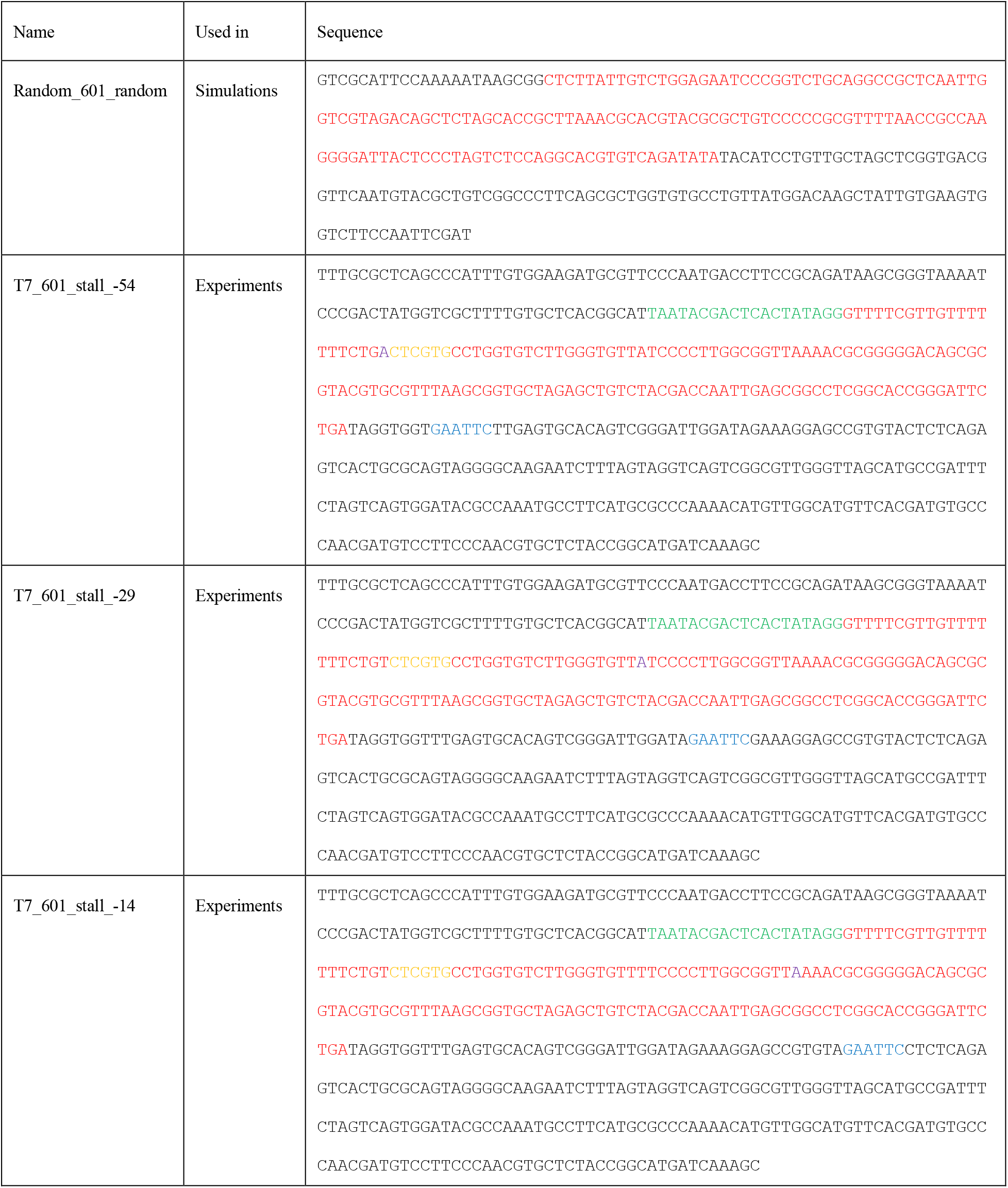
The DNA sequences used in simulations or experiments. The (modified) 601 sequence, the T7 promoter sequence, the BssSI restriction site, the EcoRI restriction site, the T7 polymerase stall site are colored red, green, yellow, blue, and purple, respectively.

